# Contextual cues as modifiers of cTBS effects on indulgent eating

**DOI:** 10.1101/538439

**Authors:** Adrian B. Safati, Peter A. Hall

## Abstract

**Background:** Prior studies have found that continuous theta burst stimulation (cTBS) targeting the left dlPFC results in reliable increases in consumption of calorie-dense food items. However, it is not known to what extent such effects are modified by cues in the immediate eating environment. Tempting environments (i.e., those saturated with appetitive eating cues) may lead to more reliance on cognitive control networks involving the dlPFC, thereby enhancing cTBS on eating.

**Objective/Hypothesis:** The objective was to examine the extent to which cTBS effects on eating would be modified by contextual cues. It was hypothesized that cTBS effects on eating would be stronger in the presence of facilitating cues.

**Methods:** Using a between-subjects factorial design, 107 adults were randomly assigned to one of four conditions: 1) active cTBS + facilitating cues, 2) sham cTBS + facilitating cues, 3) active cTBS + inhibiting cues, 4) sham cTBS + inhibiting cues. Following stimulation participants completed a flanker paradigm and a taste test during which quantity consumed was assessed surreptitiously.

**Results:** Findings revealed a significant interaction between stimulation and cue type (F(1,102)=6.235, p=.014), such that the effects of cTBS were stronger for those in the facilitating cue condition.

**Conclusions:** The effects of cTBS on eating are strengthened in the presence of facilitating cues. Methodologically speaking, facilitating cues may be a functional prerequisite for exploring cTBS effects on eating in the laboratory. Substantively, the findings also suggest that facilitating cues in the eating environment may amplify counter-intentional food indulgence in everyday life via cognitive control failure.

## Introduction

Modulation of the left dlPFC reliably alters response to appetitive, calorie-dense food items [1,2]. Such effects are more reliable with rTMS than tDCS, and when stimulation is left-sided than right-sided [1,3,4] see also [5]. The involvement of the dlPFC in dietary self-control and the propensity toward weight gain is corroborated by functional imaging studies that also implicate the dlPFC in dietary self-control and obesity [6].

Most neuromodulation studies involving eating consider stimulation parameters more carefully than food outcome measurement—for instance the type of food product and the nature of the eating environment. However, there is both theoretical and empirical justification for considering the latter two factors, when attempting to quantify the direction and magnitude of any causal effect of the dlPFC on eating outcomes. In theory, brain systems that support cognitive control have potential to be more consequential for food consumption when the food is calorie-dense than otherwise, and when environmental cues impel indulgence rather than restraint [7–9].

For instance, incentive salience of foods tend to be stronger when homeostatic feeding systems are primed by ghrelin [6,10]. Likewise, meta analytic studies have found reliable associations between cue reactivity and eating outcomes, with visual cues as powerful as the presence of real food [11–14]. For this reason, the presence of food cues in the contextual environment should amplify the causal influence of fronto-parietal control systems on eating behavior.

Several prior studies have found evidence that individual differences in Stroop performance predict actual consumption more so in the presence of facilitative visual cues than in the presence of restraint cues [15,16]. However, to date, no experimental study has examined the potential for contextual cues to moderate the impact of dlPFC function on eating in a fully factorial experiment, crossing dlPFC modulation with cue type. The present experiment is an attempt to do this using continuous theta burst stimulation to attenuate the excitability of neuron populations in the left dlPFC and observe the effect on eating in the context of randomly assigned inhibitory versus facilitative visual cues in the eating environment.

Continuous theta burst stimulation (cTBS; [17–19]) is a highly efficient variant of rTMS that reliably reduces task performance on measures of cognitive control, particularly when targeting the left dlPFC [20]. The current study examines the joint effect of left dlPFC modulation (active vs sham) and cue type (facilitating vs. inhibiting) on calorie-dense food consumption, in order to test the hypothesis that left dlPFC attenuation will result in increased consumption more so in the presence of cues that impel indulgence than when they impel restraint.

## Methods

### Participants

A total of 107 adult participants were recruited for this study. Three participants discontinued participation, leaving an effective sample size of 104 (39 males and 65 females). All participants were right handed with a mean age of 21.9 (*SD* = 3.0; range=18-37). Participants were primarily Asian (43.3%), Caucasian (27.9%), or South Asian (14.4%). Mean body mass index (BMI) was 23.0 (*SD* = 3.6; range=16.8-35.4); the majority of the sample within the “normal” range (72.8%).

Participants were recruited over 8 months (January through August, 2018) through posters distributed around the university campus. All participants were naïve to TMS; prior to participation, individuals were screened to be free of any physical and neurological conditions that would contraindicate rTMS, using a standard screening form [21]. The study protocol was reviewed by and received clearance from the institutional ethics review board. Written and informed consent was obtained from all participants prior and following to their participation. One participant discontinued participant due to reluctance to remove a head scarf for religious reasons and two discontinued due to discomfort during the motor threshold establishment procedure. In the latter two participants discomfort was alleviated immediately by discontinuing stimulation.

### Procedures

Participants were randomly assigned to one of 4 conditions: 1) active cTBS + facilitating cues, 2) sham cTBS + facilitating cues, 3) active cTBS + inhibiting cues, or 4) sham cTBS + inhibiting cues. All participants were blinded to stimulation condition. Each study session was conducted 11:00am-12:30pm or 3:00pm-4:30pm from Monday-Friday. Participants were asked to refrain from eating or consuming caffeinated beverages 3 hours prior to the start of the experimental session; adherence to these requirements was checked with completion of the consent form. All computer tasks were presented using Inquisit software version 5.0.13.0 (Millisecond Software) on a 27-inch monitor. For the cognitive tasks, participants were instructed to respond as quickly and accurately as possible. The ambient lighting and temperature conditions were maintained stable across participants. All analyses was conducted using SPSS V. 25 (IBM).

The experimental session started with the consent procedure, followed by a computer task (IAT), rTMS protocol (see below), two measures of attitudes in counterbalanced order (implicit and explicit), self-report measures (food cravings), and a computerized task of behavioral inhibition. Following the testing session—and approximately 30 minutes after stimulation—participants were given an opportunity to sample 5 different calorie dense snack foods under the guise of examining the relationship between brain function and taste perception. Change in weight of food from pre- to post-tasting was surreptitiously assessed in order to quantify food consumption. The mild deception about the purpose of the study was then explained in a debriefing session that followed; participants were then given the opportunity to withdraw their data as per ethical requirements, however none elected to do so. Following the disclosure of their study condition all participants in the sham condition reported being initially unaware that they were in the Sham condition during the stimulation protocol.

### Brain Stimulation Protocol

The cortical stimulation protocols were applied using a 75mm figure-8 coil (MCF-B65), with pulses generated by a MagPro (model X1000) biphasic stimulation unit (MagVenture, Alpharetta, GA, USA). Individualized resting motor thresholds (RMT) were employed to calibrate stimulation intensity vis-à-vis visible twitch of the right abductor *pollicus brevis (*APB) muscle. RMT was established as lowest intensity required to induce a discernable thumb twitch in 5/10 consecutive trials. The F3 electrode position (from the international 10-20 system) was used to locate target site for the left dlPFC. Stimulation intensity was set at 80% of RMT and consisted of a 40s continuous train of 600 pulses applied in the theta burst pattern (i.e., bursts of three stimuli at 50 HZ repeated at a 5 HZ frequency). Sham cTBS was applied using the placebo version of the same coil (MCF-P-B65 coil), again targeting F3.

### Flanker Task

A modified version of the Eriksen flanker task was employed as a measure of behavioral inhibition (Eriksen et al., 1974). Following a practice block of 32 trials, participants completed 5 blocks of 108 trials (96 noise, and 12 no noise) in a mixed block design. As per the original Eriksen paradigm, blocks consisted of 5 different noise conditions; the order of the trials was selected randomly but rotated such that over the course of the experiment every permutation was equiprobable. The target letters “H” and “K” were assigned to either the “A” or “D” keyboard key, while the target letters “S” and “C” were assigned to the other alternative; letter assignment was random for each participant but maintained across trials. For each trial participants were asked to stare at a fixation cross in the middle of the screen, and when they pressed the space bar a stimulus would appear. Participants were required to identify the target letter in the center of the array, ignoring any flanking noise letters and register a response using the corresponding keyboard key. Participants were allowed to proceed at their own pace, but were given a maximum of 1 s in which to respond to any given stimulus. The Flanker interference score was calculated as the difference between reaction times on correct trials in noise condition 3 (incongruent noise) and noise condition 1 (congruent noise); this served as the primary metric for subsequent analyses in the current study.

### Food Consumption

Participants were seated in front of an array of 5 snack foods, all of which were calorie-dense (2 types of salted potato chips and 3 types of Belgian chocolate balls). Participants were given a series of self-report scales to indicate the extent to which each elicited a different sensory experience (sweet, savory, etc.). The form of the taste test is commonly used in the eating literature and has been demonstrated to be a reliable metric for food consumption. Prior validation studies have shown variability in this kind of paradigm to be responsive to food palatability and level of hunger [22], and responsive to acute manipulations of executive function using cTBS targeting the left dlPFC [3,23]. Participants were given condition specific instructions during the lead-in to the taste test: participants in the facilitation condition were instructed to “eat as much as you like in order to make your ratings” while those in the inhibition condition were instructed to “eat the bare minimum in order to make your ratings.”

### Visual Cues

Participants were exposed to a visual cue containing an image of a calorie-dense food item (i.e., a pepperoni pizza) or a health-oriented informational image of the same size and shape (i.e., a circular food recommendation pyramid; ***Figure 1***). Each poster was 60*cm* x 85*cm*, and was placed on the wall at a 45 degree diagonal from the computer screen. The poster was switched for each participant in accordance with their randomly assigned cue condition. Visual cue posters were intended to be peripheral but within the visual field of each participant during the first phase of the study (e.g., consent, self-report questionnaires, and cognitive testing).

**Figure 1.**
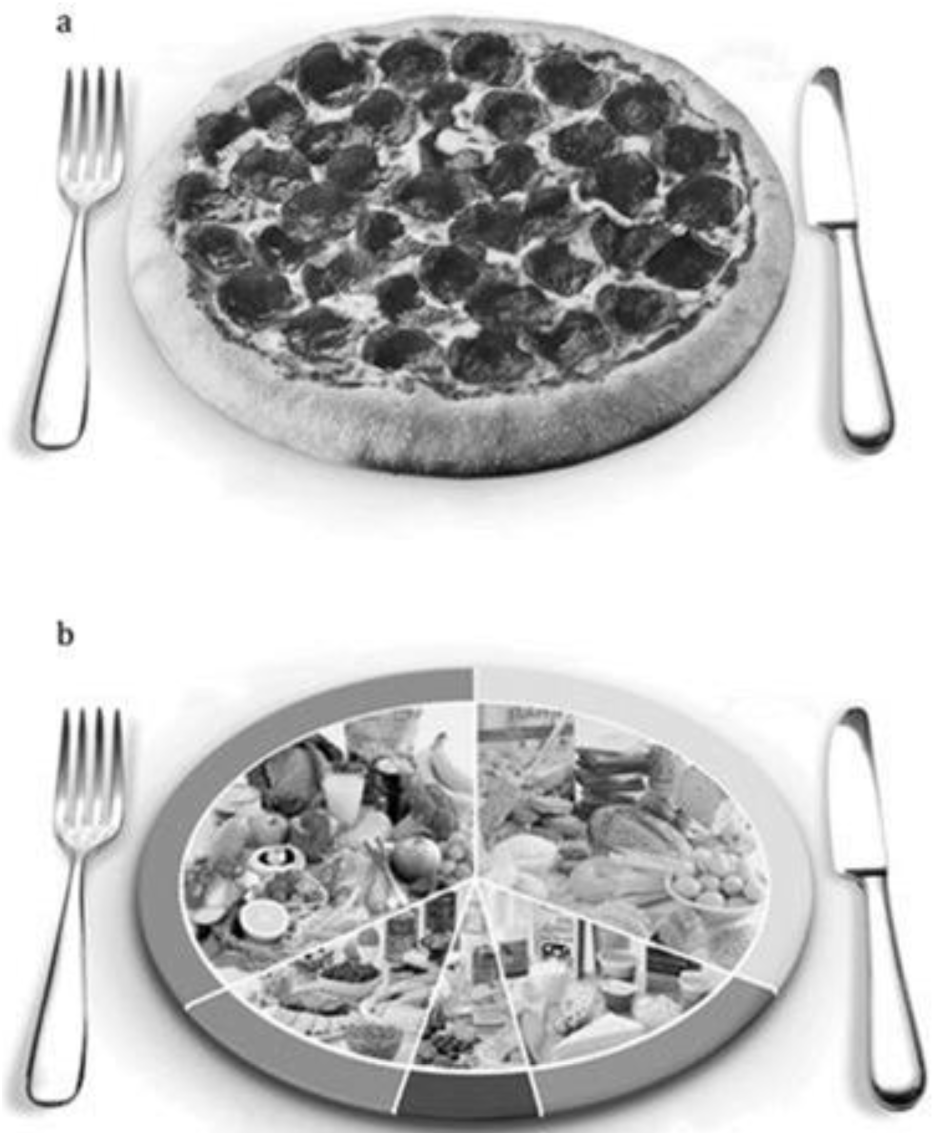
The facilitating cue poster (a) and the inhibiting cue poster (b)

### Implicit Attitudes

The IAT [24] was used to measure implicit associations between calorie density (high vs. low) and semantic valence (positive vs. negative); it was administered pre- and post-stimulation. The target food items and words were selected based on their usage in prior food IAT research [25]. Based on prior evaluative ratings of words in a large normative sample [26], the average valence of words chosen as positive words for this version of the IAT were significantly more positive than those chosen as the negative words (*t*(1,10)=7.229, *p*<.001). As in the original Greenwald study [24], the IAT consisted of 7 blocks of sorting trials. In every trial a word stimulus would be presented in the middle of the screen and participants would be required to sort it into the appropriate category on either the left or right side of the screen using the “A” key or the “D” key on the keyboard respectively. Following training blocks in which participants were required to correctly sort words according to a single category (i.e. high-calorie vs. low-calorie, or unpleasant vs. pleasant) the categories were combined (i.e. high-calorie/pleasant vs. low-calorie/unpleasant, or low-calorie/pleasant vs. high-calorie/unpleasant). The presentation order of the combined categories was randomized between participants. The primary outcome measure was a change in the D-score between the pre- and post-stimulation administrations of the IAT. The D-score was calculated as the difference in the mean sorting response times between the different combinations of category groupings (i.e. the “high-calorie/pleasant vs. low-calorie/unpleasant” blocks and the “low-calorie/pleasant vs. high-calorie/unpleasant” blocks), divided by the inclusive standard deviation of the response times in those blocks. Reaction times for trials that were more than 2 *SD* from the mean of a participant’s response times were excluded from analyses. Higher scores on the D’ metric is interpreted as a stronger positive association between high calorie foods and positive valance words.

### Food Cravings

The Food Cravings Questionnaire-State (FCQS; [27]) is a 15-item scale assessing the strength of current subjective food cravings. Higher scores on the FCQS indicate stronger craving responses experienced in the here and now. The scale includes items pertaining to the desire to eat, anticipated positive reinforcement from eating, anticipated negative reinforcement from eating, subjective lack of control over eating, and physiological symptoms of hunger.

### Explicit Attitudes

Explicit attitudes were measured using self-report. Participants were asked to rate indulgent eating using 16 sets of bipolar adjective pairs in relation to a common word stem (i.e., “for me to eat calorie dense foods would be …” wise/foolish; good/bad; etc), using a 1 to 7 scale. Responses were summed such that higher scores indicated more positive explicit attitudes toward food indulgence. This scale was previously validated and employed in an eating studies involving neuromodulation in our laboratory [28].

## Results

No significant differences were evident among the four treatment conditions with respect to age (*F*(3,103)=.136, *p*=.938), gender (*χ*(3) = 1.171, *p* = .760), BMI (*F*(3,102)=1.701, *p*=.172), time of last meal (*F*(3,103)=.561, *p*=.642), or cTBS intensity (*F*(3,103)=1.375, *p*=.255), see ***Table 1*** for details.

**Table 1.**
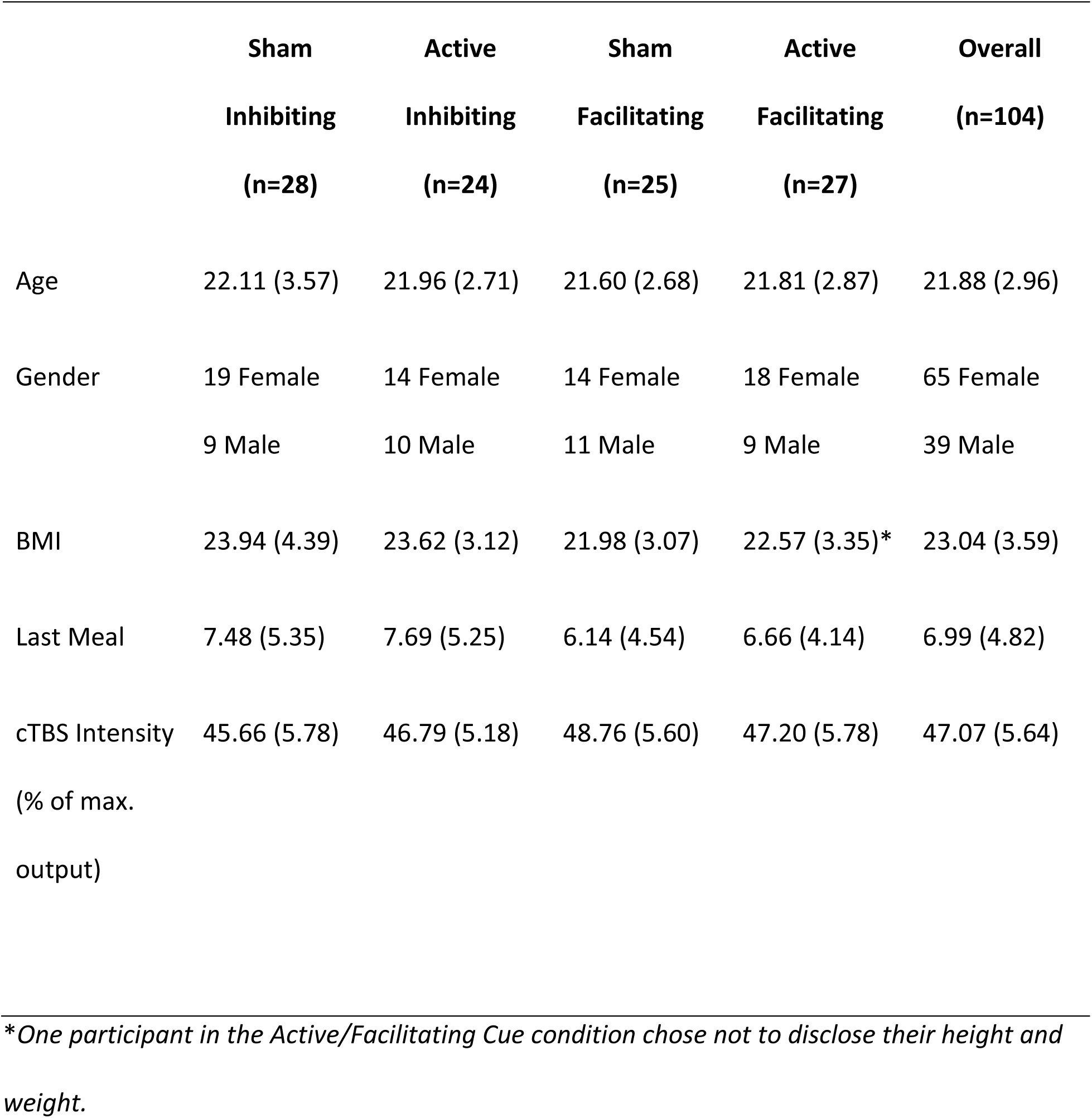
Mean (*SD*) for demographic variables by treatment condition

A univariate generalized linear model was constructed to examine the effects of stimulation condition and cue type using a two-way ANOVA for each of the outcome measures (Flanker interference scores, food consumption, cravings, explicit attitudes, and IAT scores). Gender, BMI, and dieting or sports participation were included as covariates in all analyses.

### Flanker Interference Scores

With respect to interference scores, a 2-way (stimulation condition x cue type) ANOVA revealed no main effect of cue type (*F*(1,102)=.008, *p*=.931, *g*=.017), but a significant main effect of stimulation condition (*F*(1,102)=8.844, *p*=.004, *g*=-.585), such that those in the active stimulation condition (*M*=40.446, *SE*=3.772) exhibited a stronger interference effect than those in the sham stimulation condition (*M*=24.728, *SE*=3.666). The interaction term between stimulation condition and cue type was not significant (*F*(1,102)=.001, *p*=.976). Variable means for all study conditions are depicted in ***Figure 2***.

**Figure 2.**
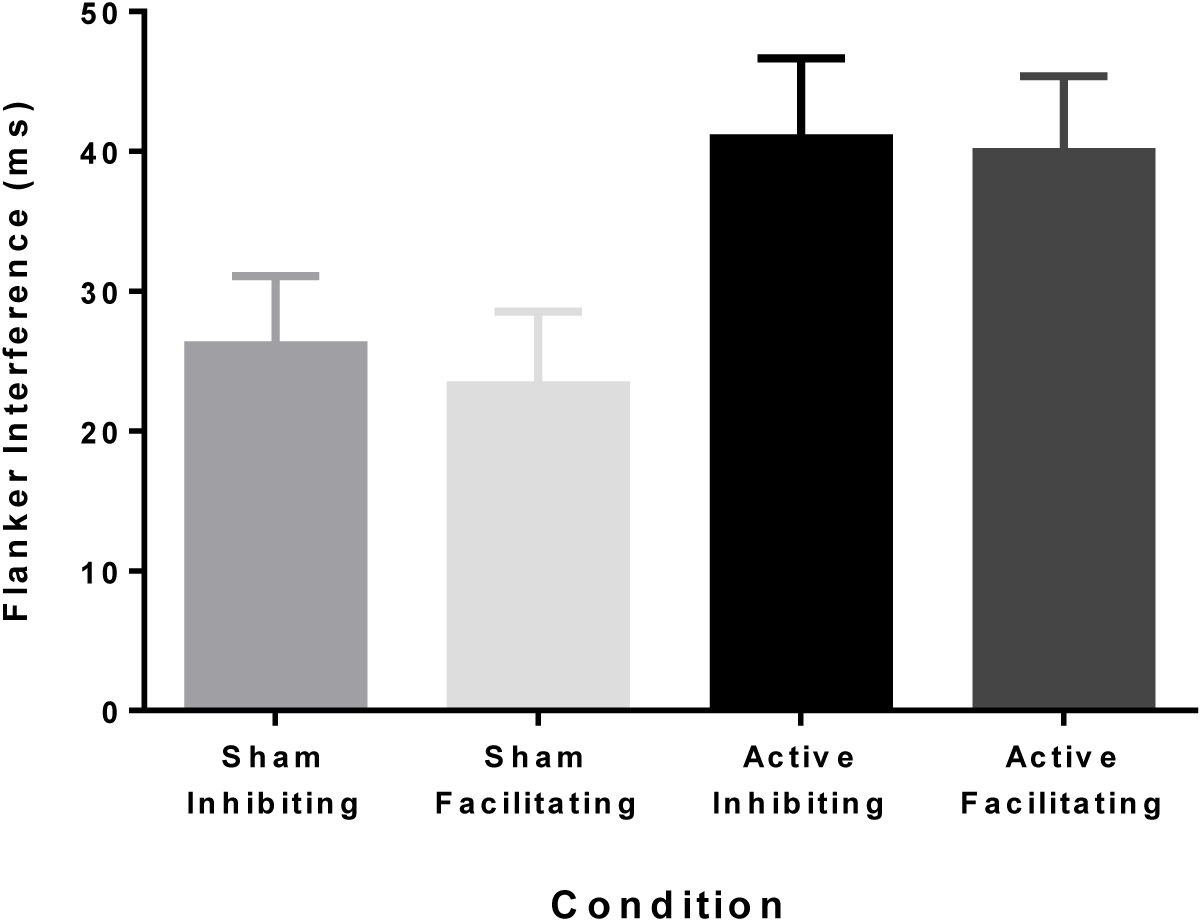
Mean (standard error) for flanker interference scores (ms) by treatment condition

### Snack Food Consumption

With respect to snack food consumption, a main effect of cue type (*F*(1,102)=15.067, *p*<.001, *g*=-0.771), was evident such that individuals in the facilitating cue conditions (*M*=79.985, *SD*=3.919) consumed significantly more snack foods than those in the inhibiting cue conditions (*M*=58.222, *SD*=3.890). There was no significant main effect of stimulation (*F*(1,102)=1.029, *p*=.313, *g*=-0.199). The effect of cue type on eating was qualified by a significant two-way interaction (*F*(1,102)=6.235, *p*=.014); means are depicted in ***Figure 3***. Consumption was greatest among those in the active condition who were exposed to facilitating cues (*M*=89.659, *SD*=5.422). Planned comparisons indicated that the difference between active and sham stimulation within the facilitation cue condition was significant (t(1,50)=12.630, p<.001) as was the difference between the cue type within the active condition (t(1,49)=4.509, p<.001).

**Figure 3.**
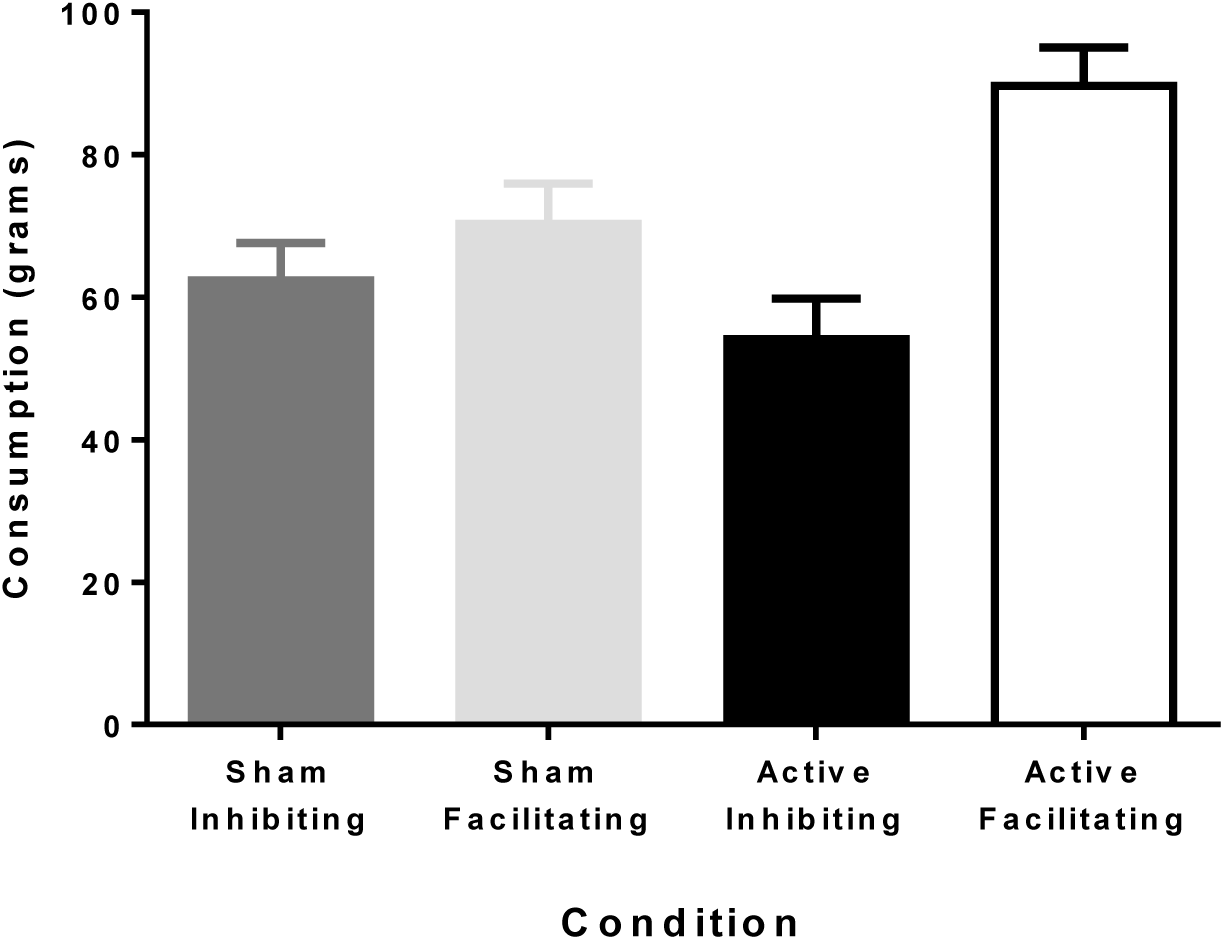
Mean (standard error) for taste test consumption (grams) by treatment condition

### Food Cravings

The two way ANOVA examining stimulation condition x cue type revealed a main effect of cue (*F*(1,102)=8.762, *p*=.004, *g*= 0.588), such that individuals in the inhibiting cue conditions (*M*=49.714, *SD*=1.355) reported increased cravings for high calorie foods compared to those in the facilitating conditions (*M*=43.934, *SD*=3.890). There was no significant main effect of stimulation (F(1,102)=1.134, p=.290, g= 0.209). The effect of cue type on eating was qualified by a two-way interaction (*F*(1,102)=8.718, *p*=.004). Means are presented in ***Figure 4***, greater values indicate stronger cravings.

**Figure 4.**
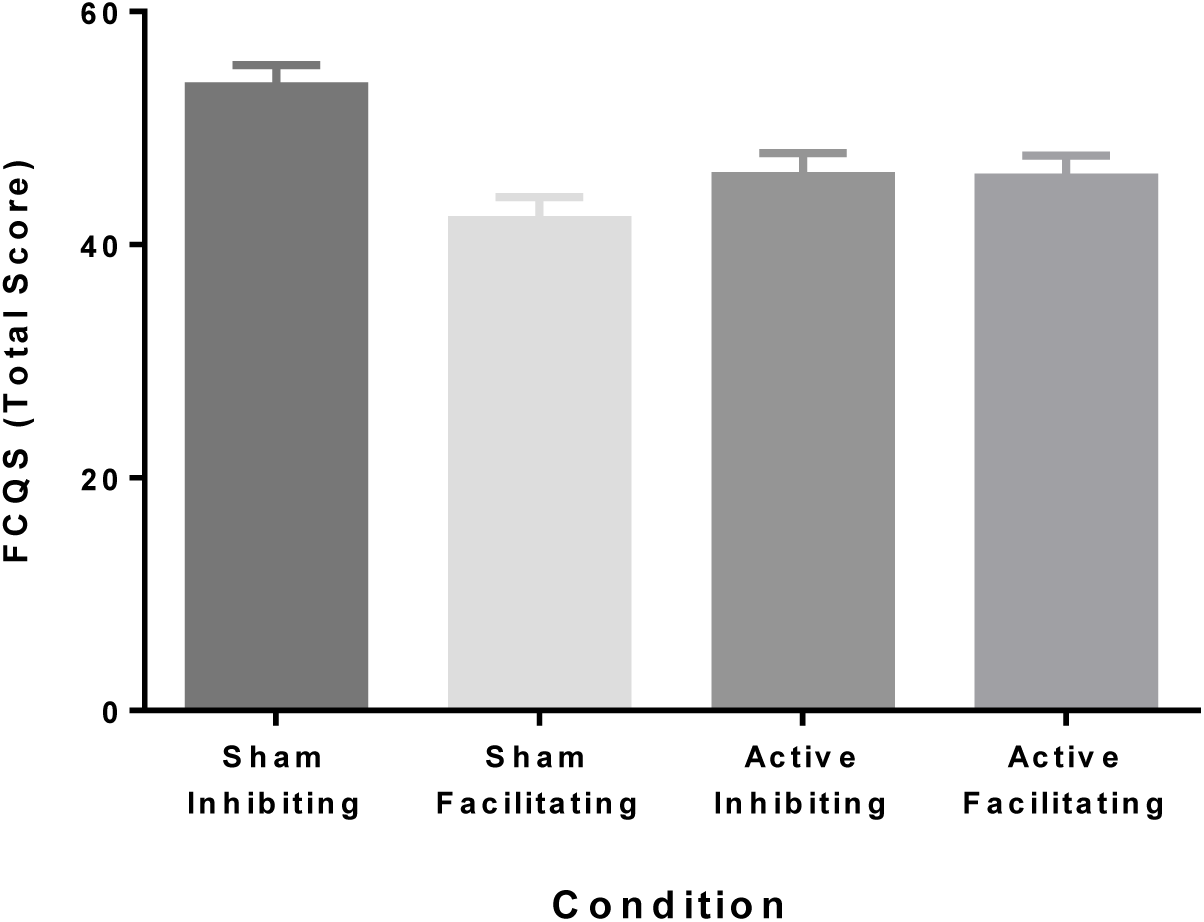
Mean (standard error) for FCQS total scores by treatment condition

### Explicit Attitudes

Two way (stimulation x cue type) ANOVA revealed no main effects of cue type (*F*(1,102)=.934, *p*=.336, *g*=0.061), or stimulation condition (*F*(1,102)=.057, *p*=.812, *g*=0.015) on explicit attitudes towards high calorie foods. The interaction term between stimulation condition and cue type was also not significant (*F*(1,102)=3.100, *p*=.081). Means are presented in ***Figure 5***. Likewise, no significant main effects or interactions emerged involving explicit attitudes toward indulgent eating.

**Figure 5.**
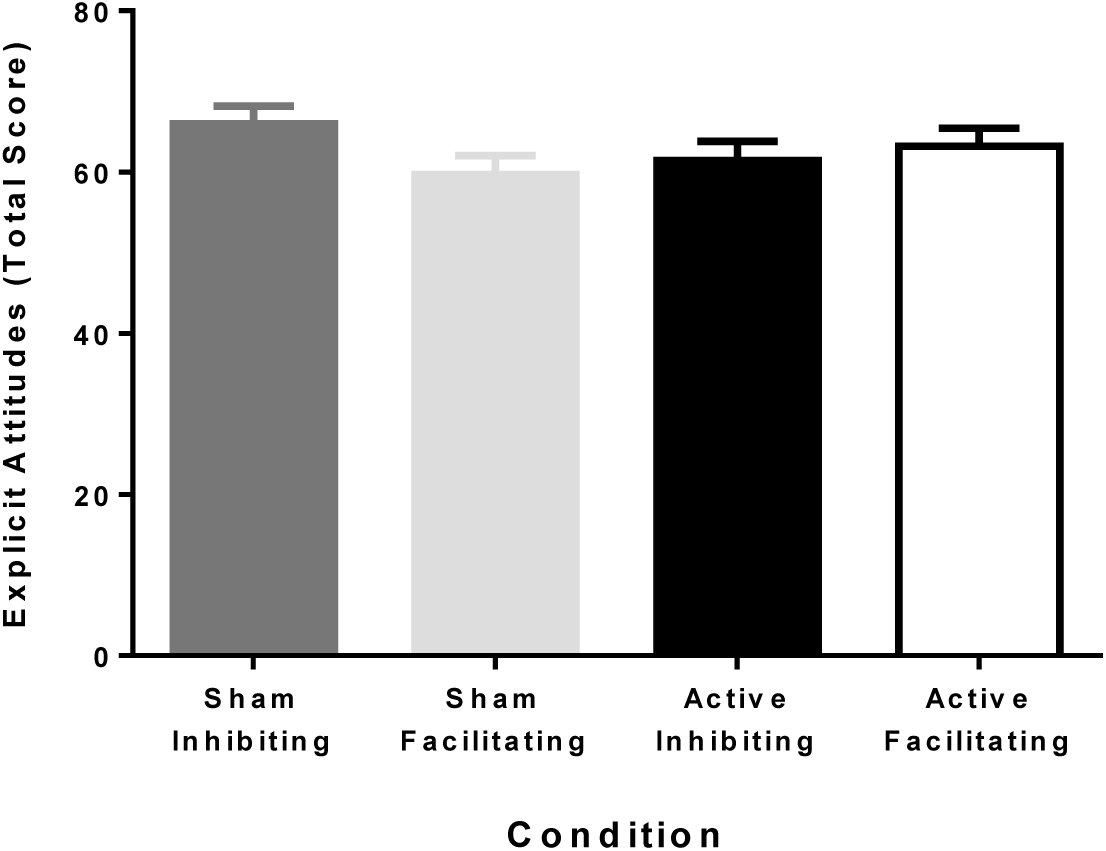
Mean (standard error) for Explicit Attitudes questionnaire total scores by treatment condition

### IAT Scores

Two way (stimulation x cue type) ANOVA revealed no main effects of cue (*F*(1,102)=.036, *p*=.850, *g*=0.039), or stimulation (*F*(1,102)=3.149, *p*=.079, *g*=0.353) on a change in implicit attitudes. The interaction term between stimulation and environment was not significant (*F*(1,102)=1.224, *p*=.271). Means are presented in ***Figure 6.***

**Figure 6.**
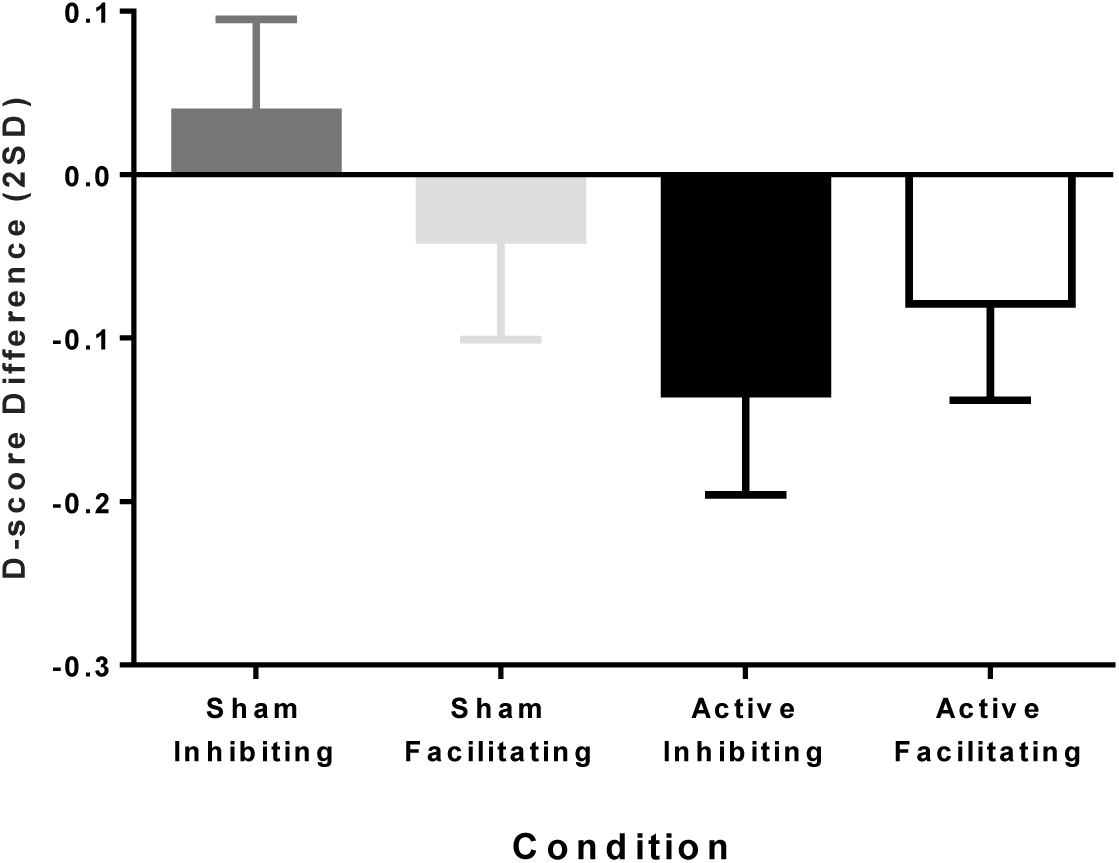
Mean (standard error) for changes in D-score in IAT performance pre- and post-stimulation by treatment condition

## Discussion

The current study employed a between-subjects factorial to design to test the hypothesis that the left dlPFC modulation of eating behavior would be more apparent when cues were facilitative of indulgence than otherwise. Using cTBS to modulate the target site, we found that active stimulation (i.e., attenuation of excitability of neuron populations in the left dlPFC) resulted in significantly more eating when environmental cues were facilitative than when they were inhibiting.

Our findings augment existing experimental neuromodulation research involving eating by identifying an important contextual parameter of the eating environment that may determine the magnitude of experimental effect to be expected from neuromodulation. Variability in findings of dlPFC modulation and eating outcome in the existing research literature [1,4] could potentially be explained by unintended variability in the eating environment and the extent to which available cues impel restraint or indulgence, even indirectly.

Although it is the case that our visual cue manipulation was one that was more clear than others, it is possible that more subtle cues could result in the same effects. For instance, an experimental setting that contains incidentally visible food images that are appetizing might introduce similar effects; likewise, situations wherein participants are presented with large numbers of appetitive food images prior to the food opportunity for task purposes might do the same. Conversely, studies without such cues and tasks prior to the eating opportunity might find smaller effects of stimulation. Studies executed in clinical settings might inadvertently prime health consciousness and reluctance to indulge, thereby reducing effect size of control network modulation on eating outcomes.

The current findings also have substantive meaning beyond the methodological implications. Given that advertising for food items in the modern living environment rely on appetizing images, it is possible that such advertising may result in acute susceptibility to indulgence, particularly when other acute dlPFC suppressing factors are present, such as sleep deprivation [29], stress [30], or alcohol intoxication [31]. From this perspective, the key to increasing probability of successfully resisting indulgence relies partially on avoiding exposure to acute attenuators of cognitive control such as the above.

Strengths of this study include: 1) the large sample size, allowing for between condition comparisons that would not be sufficiently powered in more conventionally sized neuromodulation study samples, 2) the employ of a between-subjects design, which enhances the validity of the findings by reducing the chance of loss of blinding, and finally 3) the use of sham coil which, in conjunction with the between-subjects design, further enhances the validity of the experimental conditions and reduces the ability of participants to compare stimulation sensations across conditions.

In conclusion, the current study found evidence that the effect of dlPFC attenuation via cTBS is stronger under conditions of behavior-facilitative cues. Findings suggest that neuromodulation studies involving eating should include appetitive cues in the eating environment and/or avoid incidental exposure to inhibiting cues. Perhaps even more important are the implications of the current findings for when self-restraint would be expected to be more taxing of cognitive control networks in everyday life.

## Acknowledgements

Support for this work was provided in part by an operating grant to the second author from the Social Sciences and Humanities Research Council of Canada (435-2017-0027).

## References

[1] Lowe CJ, Vincent C, Hall PA. Effects of Noninvasive Brain Stimulation on Food Cravings and Consumption: A Meta-Analytic Review. Psychosom Med 2017;79:2–13. doi:10.1097/PSY.0000000000000368.

[2] Hall PA, Lowe CJ. Cravings, currents and cadavers: What is the magnitude of tDCS effects on food craving outcomes? Nutr Neurosci 2018:1–4. doi:10.1080/1028415X.2018.1513678.

[3] Lowe CJ, Hall PA, Staines WR. The effects of continuous theta burst stimulation to the left dorsolateral prefrontal cortex on executive function, food cravings, and snack food consumption. Psychosom Med 2014;76:503–11. doi:10.1097/PSY.0000000000000090.

[4] Hall PA, Lowe C, Vincent C. Brain Stimulation Effects on Food Cravings and Consumption: An Update on Lowe et al. (2017) and a Response to Generoso et al. (2017). Psychosom Med 2017;79:839–42. doi:10.1097/PSY.0000000000000504.

[5] Kim S-H, Chung J-H, Kim T-H, Lim SH, Kim Y, Lee Y-A, et al. The effects of repetitive transcranial magnetic stimulation on eating behaviors and body weight in obesity: A randomized controlled study. Brain Stimulat 2018;11:528–35. doi:10.1016/j.brs.2017.11.020.

[6] Han JE, Boachie N, Garcia-Garcia I, Michaud A, Dagher A. Neural correlates of dietary self-control in healthy adults: A meta-analysis of functional brain imaging studies. Physiol Behav 2018;192:98–108. doi:10.1016/j.physbeh.2018.02.037.

[7] Hall PA. Executive-Control Processes in High-Calorie Food Consumption. Curr Dir Psychol Sci 2016;25:91–8. doi:10.1177/0963721415625049.

[8] Hall PA, Bickel WK, Erickson KI, Wagner DD. Neuroimaging, neuromodulation, and population health: the neuroscience of chronic disease prevention. Ann N Y Acad Sci 2018;1428:240–56. doi:10.1111/nyas.13868.

[9] Tang DW, Fellows LK, Dagher A. Behavioral and neural valuation of foods is driven by implicit knowledge of caloric content. Psychol Sci 2014;25:2168–76. doi:10.1177/0956797614552081.

[10] Han JE, Frasnelli J, Zeighami Y, Larcher K, Boyle J, McConnell T, et al. Ghrelin Enhances Food Odor Conditioning in Healthy Humans: An fMRI Study. Cell Rep 2018;25:2643–2652.e4. doi:10.1016/j.celrep.2018.11.026.

[11] Boswell RG, Kober H. Food cue reactivity and craving predict eating and weight gain: a meta-analytic review. Obes Rev Off J Int Assoc Study Obes 2016;17:159–77. doi:10.1111/obr.12354.

[12] Lawrence NS, Hinton EC, Parkinson JA, Lawrence AD. Nucleus accumbens response to food cues predicts subsequent snack consumption in women and increased body mass index in those with reduced self-control. NeuroImage 2012;63:415–22. doi:10.1016/j.neuroimage.2012.06.070.

[13] Lopez RB, Hofmann W, Wagner DD, Kelley WM, Heatherton TF. Neural predictors of giving in to temptation in daily life. Psychol Sci 2014;25:1337–44. doi:10.1177/0956797614531492.

[14] Yokum S, Gearhardt AN, Harris JL, Brownell KD, Stice E. Individual differences in striatum activity to food commercials predict weight gain in adolescents. Obes Silver Spring Md 2014;22:2544–51. doi:10.1002/oby.20882.

[15] Hall P, Tran B, Lowe C, Vincent C, Mourtzakis M, Liu-Ambrose T, et al. Expression of executive control in situational context: Effects of facilitating versus restraining cues on snack food consumption. Health Psychol Off J Div Health Psychol Am Psychol Assoc 2015;34:539–46. doi:10.1037/hea0000134.

[16] Hall PA, Lowe C, Vincent C. Executive control resources and snack food consumption in the presence of restraining versus facilitating cues. J Behav Med 2014;37:587–94. doi:10.1007/s10865-013-9528-3.

[17] Huang Y-Z, Edwards MJ, Rounis E, Bhatia KP, Rothwell JC. Theta burst stimulation of the human motor cortex. Neuron 2005;45:201–6. doi:10.1016/j.neuron.2004.12.033.

[18] Huang Y-Z, Rothwell JC, Chen R-S, Lu C-S, Chuang W-L. The theoretical model of theta burst form of repetitive transcranial magnetic stimulation. Clin Neurophysiol Off J Int Fed Clin Neurophysiol 2011;122:1011–8. doi:10.1016/j.clinph.2010.08.016.

[19] Suppa A, Huang Y-Z, Funke K, Ridding MC, Cheeran B, Di Lazzaro V, et al. Ten Years of Theta Burst Stimulation in Humans: Established Knowledge, Unknowns and Prospects. Brain Stimulat 2016;9:323–35. doi:10.1016/j.brs.2016.01.006.

[20] Lowe CJ, Manocchio F, Safati AB, Hall PA. The effects of theta burst stimulation (TBS) targeting the prefrontal cortex on executive functioning: A systematic review and meta-analysis. Neuropsychologia 2018;111:344–59. doi:10.1016/j.neuropsychologia.2018.02.004.

[21] Rossi S, Hallett M, Rossini PM, Pascual-Leone A, Safety of TMS Consensus Group. Safety, ethical considerations, and application guidelines for the use of transcranial magnetic stimulation in clinical practice and research. Clin Neurophysiol Off J Int Fed Clin Neurophysiol 2009;120:2008–39. doi:10.1016/j.clinph.2009.08.016.

[22] Robinson E, Haynes A, Hardman CA, Kemps E, Higgs S, Jones A. The bogus taste test: Validity as a measure of laboratory food intake. Appetite 2017;116:223–31. doi:10.1016/j.appet.2017.05.002.

[23] Lowe CJ, Staines WR, Manocchio F, Hall PA. The neurocognitive mechanisms underlying food cravings and snack food consumption. A combined continuous theta burst stimulation (cTBS) and EEG study. NeuroImage 2018;177:45–58. doi:10.1016/j.neuroimage.2018.05.013.

[24] Greenwald A., McGhee D., Schwartz JL. Measuring individual differences in implicit cognition: The implicit association test. Journal of Personality and Social Psychology 1998;74:1464–80. doi:10.1037/0022-3514.74.6.1464.

[25] Mattavelli G, Zuglian P, Dabroi E, Gaslini G, Clerici M, Papagno C. Transcranial magnetic stimulation of medial prefrontal cortex modulates implicit attitudes towards food. Appetite 2015;89:70–6. doi:10.1016/j.appet.2015.01.014.

[26] Warriner AB, Kuperman V, Brysbaert M. Norms of valence, arousal, and dominance for 13,915 English lemmas. Behav Res Methods 2013;45:1191–207. doi:10.3758/s13428-012-0314-x.

[27] Cepeda-Benito A, Gleaves DH, Williams TL, Erath SA. The development and validation of the state and trait food-cravings questionnaires. Behav Ther 2000;31:151–73. doi:10.1016/S0005-7894(00)80009-X.

[28] Hall PA, Vincent CM, Burhan AM. Non-invasive brain stimulation for food cravings, consumption, and disorders of eating: A review of methods, findings and controversies. Appetite 2018;124:78–88. doi:10.1016/j.appet.2017.03.006.

[29] Lowe CJ, Safati A, Hall PA. The neurocognitive consequences of sleep restriction: A meta-analytic review. Neurosci Biobehav Rev 2017;80:586–604. doi:10.1016/j.neubiorev.2017.07.010.

[30] Qin S, Hermans EJ, van Marle HJF, Luo J, Fernández G. Acute psychological stress reduces working memory-related activity in the dorsolateral prefrontal cortex. Biol Psychiatry 2009;66:25–32. doi:10.1016/j.biopsych.2009.03.006.

[31] Abernathy K, Chandler LJ, Woodward JJ. Alcohol and the prefrontal cortex. Int Rev Neurobiol 2010;91:289–320. doi:10.1016/S0074-7742(10)91009-X.

